# Quantitative chromatin protein dynamics during replication origin firing in human cells

**DOI:** 10.1101/2024.04.25.590934

**Authors:** Sampath Amitash Gadi, Ivo Alexander Hendriks, Christian Friberg Nielsen, Petya Popova, Ian D. Hickson, Michael Lund Nielsen, Luis Toledo

**Author notes:** Current address: Protein Signalling Program, Novo Nordisk Foundation Center for Protein Research, Faculty of Health and Medical Sciences, University of Copenhagen, Copenhagen 2200, Denmark (Sampath Amitash Gadi). Current address: BiOrigin, Ole Maaløes Vej 3, Copenhagen 2200, Denmark (Luis Toledo). These authors contributed equally (Sampath Amitash Gadi, Ivo Alexander Hendriks).

## Abstract

Accurate genome duplication requires a tightly regulated DNA replication program, which relies on the fine regulation of origin firing. While the molecular steps involved in origin firing have been determined predominantly in budding yeast, the complexity of this process in human cells has yet to be fully elucidated. Here, we describe a straightforward proteomics approach to systematically analyse protein recruitment to the chromatin during induced origin firing in human cells. Using a specific inhibitor against CHK1 kinase, we induced a synchronised wave of dormant origin firing (DOF) and assessed the S phase chromatin proteome at different time points. We provide time-resolved loading dynamics of 3,269 proteins, including the core replication machinery and origin firing factors. This dataset accurately represents known temporal dynamics of proteins on the chromatin during the activation of replication forks and the subsequent DNA damage due to the hyperactivation of excessive replication forks. Finally, we used our dataset to identify the condensin II subunit NCAPH2 as a novel factor required for efficient origin firing and replication. Overall, we provide a comprehensive resource to interrogate the protein recruitment dynamics of replication origin firing events in human cells.

## Introduction

The human genome is replicated from thousands of replication origins simultaneously. These origins are licensed in G1 phase and are activated (‘fired’) in S phase. Origin firing (OF) requires the regulated recruitment of several proteins to origins of replication. The minimal set of proteins required for origin firing was established by reconstituting the entire process using purified proteins in budding yeast ^1^ and, recently, in human ^2^. These proteins are thought to be largely conserved in higher eukaryotes, including humans. First, inactive MCM helicase complexes are loaded onto DNA during licensing in G1. These inactive MCMs at the origins are also called pre-replication complexes (pre-RCs) ^3^. In S phase, the DBF4-dependent CDC7 (DDK) and CDK2 kinases activate the pre-RCs, leading to the recruitment of CDC45 and GINS to MCMs, forming the pre-initiation complexes (pre-ICs). DDK-mediated phosphorylation of MCMs at the pre-RC acts as a scaffold for the recruitment of TRESLIN-MTBP and RECQL4, although recently RECQL4 has been shown to be dispensable for OF ^4^. CDK2-mediated phosphorylation of TRESLIN leads to the formation of the TRESLIN-MTBP-TOPBP1 complex to promote CDC45, GINS and POLE recruitment. Pre-ICs give rise to two active CMG complexes (CDC45-MCM-GINS), which will form the two outgoing replisomes ^5–9^. Furthermore, MCM10 is required to initiate the unwinding of the DNA by the CMG helicase ^10^. Together, these proteins constitute a set of initiation factors required for the initial steps of origin firing ^1,11,12^.

OF leads to the formation of bi-directional replication forks, one for each CMG complex. At each fork, additional accessory proteins are recruited to the CMG to complete the formation of the replisome ^13^. These include PCNA ^14^, lagging strand polymerase δ (POLD) ^15^, the RFC complex (RFC1-5) ^16^, the RPA complex (RPA1-3) ^17^, the primase – polymerase α complex (POLA), CTF4 ^18^, the fork protection complex (FPC) ^19^ and the alternative clamp loading complex comprising CTF8, CTF18 and DCC1 ^16,20^. OF is regulated through fluctuations in CDK2 activity mediated by the S phase checkpoint kinases ATR and CHK1. Inhibition of these kinases through specific inhibitors (ATRi and CHK1i) induces dormant origin firing (DOF) ^17,21^.

While much of our knowledge of OF has been derived from functional studies in budding yeast and in cell-free extracts from *Xenopus* eggs, a systematic study of the origin firing process in human cells is still missing ^22^. Development of techniques such as isolation of proteins on nascent DNA (iPOND) ^23^ and nascent chromatin capture (NCC) ^23,24^ have helped to isolate factors with roles in DNA replication. These techniques, coupled with mass spectrometry (MS), enabled a discovery approach to identify factors recruited to nascent DNA under various conditions ^24–28^. However, by capturing proteins on nascent DNA, these techniques do not allow probing for proteins recruited to origins before the activation of replication forks.

Here, using quantitative proteomics, we profiled chromatin-bound protein fractions of human cells in which we induced origin firing. Our dataset provides precise profiles of the dynamic loading of 3,269 proteins over a time-course of increasing origin firing and activation of new replication forks. We provide an extensive map of time-resolved chromatin-loading dynamics of proteins with different roles in DNA replication, including origin firing factors, proteins of the core replisome, and proteins involved in the response to DNA damage caused by RPA exhaustion. Finally, our loading dynamics analysis found several proteins with uncharacterized roles in replication and led us to identify NCAPH2 as a new factor required for origin firing in human cells.

## Results

### CHK1i induces a CDC7 and CDK-dependent wave of origin firing in S phase

OF in eukaryotes is a multi-step process leading to the activation of replication forks ^10^. First, CDC7 and CDK2 activities convert pre-RCs into pre-ICs through the recruitment of OF factors. This is followed by the unwinding of the double-stranded DNA to generate a replication bubble containing two replication forks (RFs; Figure 1A). To assess all three steps of OF (pre-RCs, pre-ICs and RFs) in human cells, we used the checkpoint inhibitor AZD7762 ^29,30^, which strongly inhibits CHK1 and to a lesser extent CHK2. However, CHK2 is largely inactive in the absence of DNA damage, whereas CHK1 is active throughout an unperturbed S phase ^31,32^. Our focus is on OF and we believe it reasonable to assume that the effect of AZD7762 on OF will be limited to inhibiting CHK1 in S phase cells. Quantitative image-based cytometry (QIBC) analysis showed that a 30-minute treatment of U2OS cells with 1 µM of CHK1i induced a strong upregulation in chromatin-bound RPA (CB-RPA) in S phase cells, which was entirely rescued upon inhibition of CDC7 or CDKs (Figure 1B). We also found a temporal increase in the chromatin-bound levels of the replication OF factors Treslin, RECQL4, TOPBP1, RPA70, CDC45, GINS3, and PCNA (Figures 1C and S1A). This was consistent with data obtained using QIBC which showed a temporal increase in CB-RPA, but no concomitant increase in γH2AX until the 30’ time point (Figures 1D, S1B and S1C). γH2AX sharply increased at the 60’ time point specifically in cells with already high CB-RPA, a characteristic feature of Replication Catastrophe (RC) (Figure S1C)^17^. To confirm the effect of CHK1i on OF, we also probed CDK2 activity by measuring phosphorylation of one of its targets, FOXM1 at site T600 (pFOXM1). We observed a temporal increase in pFOXM1 levels in cells treated with CHK1i (Figures 1E and S1D). Our QIBC analysis was able to capture an increase in pFOXM1 levels as early as 5 minutes after treatment with CHK1i. Slightly later, this was followed by an increase in CB-RPA (Figure 1F). This is consistent with the requirement of CDK2 activity upstream of replication initiation, after which RPA is loaded on to the chromatin. Altogether, our data validates that the CHK1i AZD7762 is a potent inducer of OF.

**Figure 1.**
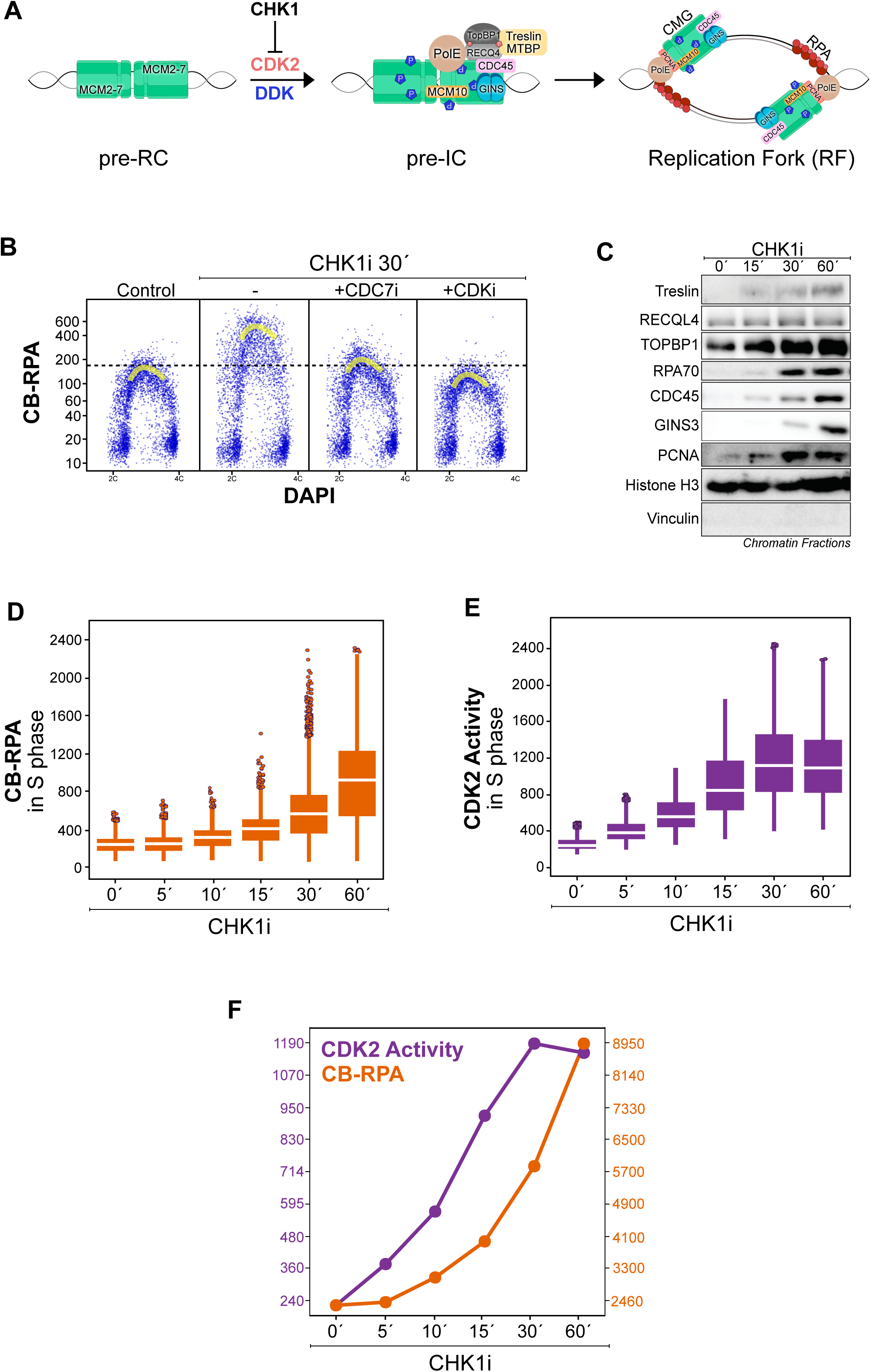
– CHK1i induces a CDC7 and CDK-dependent wave of origin firing in S phase. A. Schematic illustration of the OF process showing three crucial steps – preRC, preIC and the formation of the RF. CHK1 regulates CDK2 activity which is required, along with DDK to phosphorylate and activate OF factors. DDK phosphorylates MCMs to convert pre-RC to pre-IC. CDK2 phosphorylates TRESLIN-MTBP and RECQL4 to recruit TOPBP1 and the CMG forms the replisome. MCM10 assists in the unwinding of DNA to form the RF ^1^. RPA is bound to ssDNA generated at the RF. B. QIBC scatter plots of cells either untreated (-) or treated with CHK1i for 30 minutes with or without a CDK inhibitor or a CDC7 inhibitor. Yellow curves represent the peak of RPA in the scatter plot. C. Cells were treated for the indicated times with CHK1i inhibitor and chromatin-enriched fractions were prepared and analysed by western blotting with the indicated antibodies. H3 was used as a loading control. Vinculin was used as a negative control for chromatin. D. QIBC scatter plots of CB-RPA in S phase cells (gated for EdU positive) treated with CHK1i for the indicated times. E. QIBC scatter plots of FOXM1 pT600 (CDK2 activity) in S phase cells (gated for EdU positive) treated with CHK1i for the indicated times. F. Line diagram comparing the increase in CB-RPA (orange) vs FOXM1 pT600 (CDK2 activity; purple) from D and E.

### Proteomic analysis of the chromatin in CHK1i treated cells

We aimed to generate a complete picture of OF, including pre-RCs, pre-ICs and RFs through a temporal analysis of S phase chromatin proteome in CHK1i treated cells. For this, we developed a straightforward, label-free proteomics approach to survey for factors recruited to the chromatin (Figure 2A). First, we synchronised U2OS cells in S phase using a single thymidine block for 16h and then released into S phase for 4 hours. We consistently obtained more than 90% cells in S phase using this procedure (Figure S2A, see top panel 0’). We then treated the cells released into S phase with CHK1i in a time course for 5 to 120 minutes such that the end of the 4h release from thymidine coincided with the end of CHK1i treatment (Figures 2A and S2A, top panel time course). CB-RPA increased in a time-dependent manner, specifically in S phase cells, confirming induction of OF (Figure S2A, lower panel). At the end of the treatment, we collected these cells (5 replicates, see methods for details) from all the time-points, isolated the chromatin fraction, performed in-solution digestion of proteins, performed off-line sample fractionation, and subjected all samples to analysis by high-resolution mass spectrometry (MS, see methods for full details). Our MS analysis quantified 3269 proteins in quintuplicate across all time-points (Supplementary Table 1). Data reproducibility was exceptionally high, with Pearson correlations within conditions ranging from 0.988 to 0.990 (Figure S2B). Gene ontology (GO) term annotations categorised 130 of these factors in ‘DNA replication’ (GO:0006260); 161 factors in ‘DNA repair’ (GO:); 290 factors in ‘Chromatin organisation’ (GO:) and > 200 factors categorised as ‘Cell cycle’ (GO:), ‘Transcription’ (GO:) and ‘Other chromatin’ (GO:) (Figure 2B). While the majority of proteins quantified in the samples did not alter in level upon CHK1i treatment, principal component analysis revealed a notable variation in protein abundance at the various time-points (Figure S2C). Importantly, our temporal analysis comparing the different CHK1i treatment time-points to each other and to untreated S phase chromatin (time 0’) allowed us to generate a complete picture of factors loaded and unloaded from the chromatin after CHK1i treatment (Figure 2C). Overall, our dataset provides a comprehensive resource of the dynamics of the chromatin proteome during OF (Supplementary Table 1).

**Figure 2.**
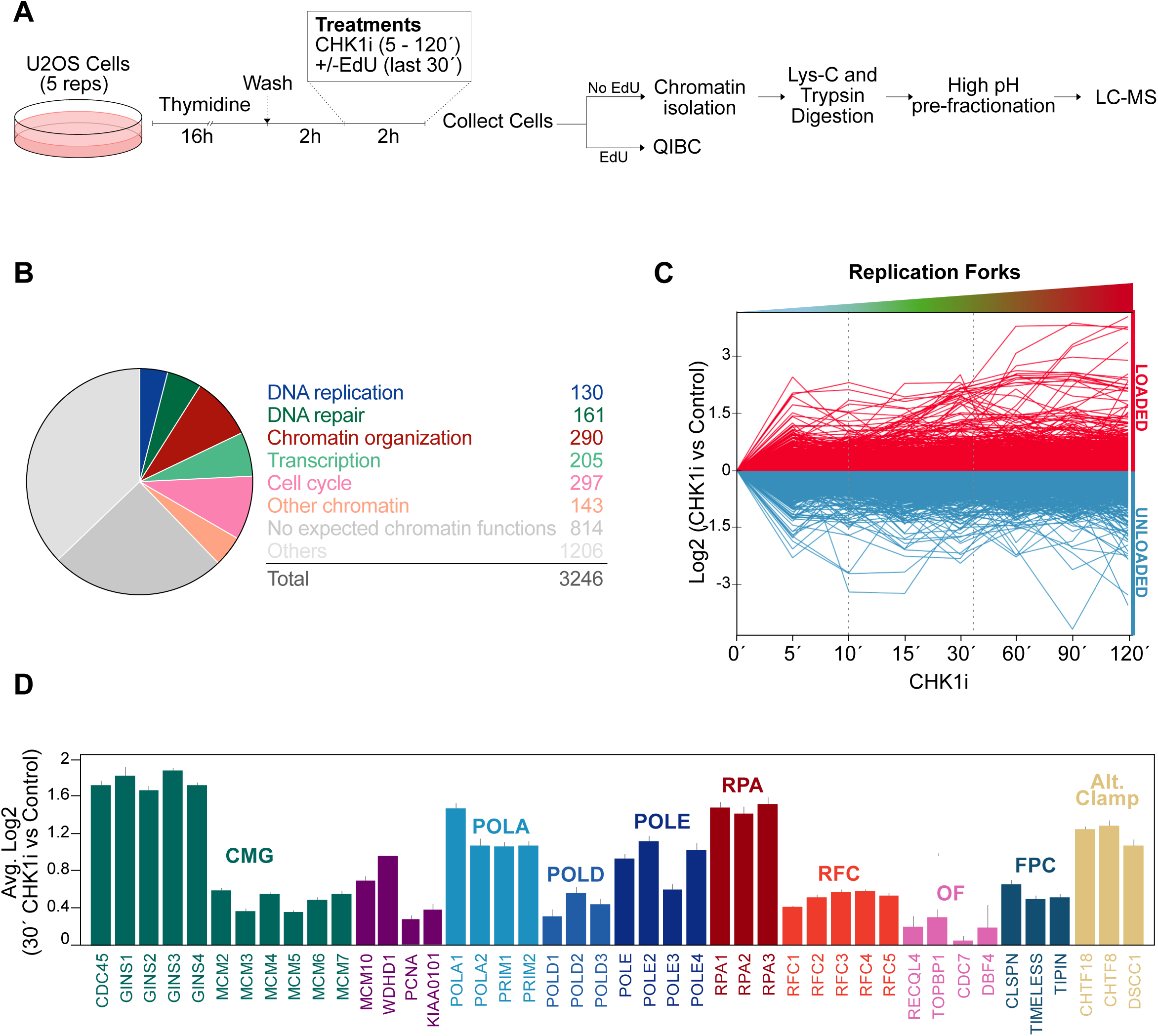
– Proteomic analysis of the chromatin in CHK1i treated cells. A. Outline of the experimental approach employed to isolate chromatin for MS and QIBC analysis. B. Pie diagram showing the distribution of all factors quantified in our MS dataset by gene ontology (GO) analysis. C. Line diagrams showing the logarithmic ratio of intensities of CHK1i treated cells in comparison with control. All 3269 quantifiable proteins from our MS analysis are displayed as either loaded (red) or unloaded (blue) on the chromatin. D. Bar graphs showing the logarithmic ratio of chromatin protein intensities of replisome factors in 30’ CHK1i treated cells compared to untreated cells. CMG (green); POLA (blue); POLD (dark blue); POLE (darker blue); RPA (red); RFC (orange); Origin Firing (OF; pink); FPC (blue grey) and the alternative clamp loader (gold) are shown.

To check the validity of our dataset, we analysed the loading of known replication factors on chromatin after treatment with CHK1i for 30’. We identified almost all known core replisome proteins, including the CMG (CDC45, GINS and MCMs), MCM10, PCNA, POLA, POLD, POLE, RPA, RFC, the DDK, the fork protection complex (FPC) and the alternative clamp loader proteins (Figure 2D). We were unable to quantify POLD4 and Treslin-MTBP. POLD4 is a very small protein (∼12 kDa), generating very few peptides and making it hard to detect by MS ^33,34^. Five unique peptides for Treslin were detected, but only in three replicates, and thus could not be quantified in the same way. Notably, these proteins are also absent in datasets from the more commonly used techniques like iPOND (isolation of proteins on nascent DNA) and NCC (nascent chromatin capture) ^24,34^. Further comparison of our dataset with iPOND and NCC datasets showed that we were able to quantify most of the missing or difficult to detect replication proteins (Figure S2D and Supplementary Table 1). Taken together, our MS methodology, which essentially profiles the whole chromatin, does not rely on convoluted labelling, crosslinking, or enrichment techniques, and yet detects proteins quantitatively by the presence of a high number of peptides. Importantly, this enabled a high degree of detection sensitivity as well as exceptional quantitative accuracy and precision.

### Time-resolved dynamics of chromatin proteins in CHK1i-treated cells

One of the key aspects of our dataset is the temporal dynamics of proteins on the chromatin when OF was induced with CHK1i. To better illustrate these dynamics, we plotted the quantified values from the entire time-course together with the control (time-point 0’) as line graphs (Figure 3). The core replicative helicase consisting of CDC45, GINS, and MCMs (CMG) showed a time-dependent increase in loading on the chromatin in CHK1i-treated cells (Figure 3A). This indicates a gradual conversion of pre-RCs into pre-ICs and the formation of RFs as more and more CMGs are assembled due to increase in CDK2 activity. Importantly, the levels of core histones, indicating total chromatin, remained unchanged throughout the time-course (Figure S3A). Similarly to the CMG, other replication initiation factors including RPA, RFC, FPC, AND-1, MCM10, and the alternative RFC complex (comprising CHTF8, CHTF18 and DSCC1) also showed a time-dependent increase in loading on the chromatin in CHK1i-treated cells (Figure S3B-E). Transient initiation factors CDC7, TOPBP1, and RECQL4, showed only a marginal increase in loading on the chromatin up to the 60’ time-point (Figure S3F). Interestingly, the replisome clamp PCNA and its associated factor KIAA0101 (PCLAF) showed an initial unloading trend for the first 10 minutes and only then began to enrich on the chromatin in CHK1i-treated cells (Figure 3B). The POLA complex, required for priming during DNA synthesis, also showed a gradual increase in loading up to 30’ after treatment with CHK1i, after which a rather abrupt drop in loading of this complex was observed (Figure 3C). We speculate that this could be due to fork breakage in RC. Altogether, we have captured time-resolved chromatin dynamics of essential replication proteins in CHK1i-treated cells. Importantly, subunits of complexes in our dataset showed near-identical chromatin dynamics, demonstrating the robustness of our method and dataset (Figures 3A-C).

**Figure 3.**
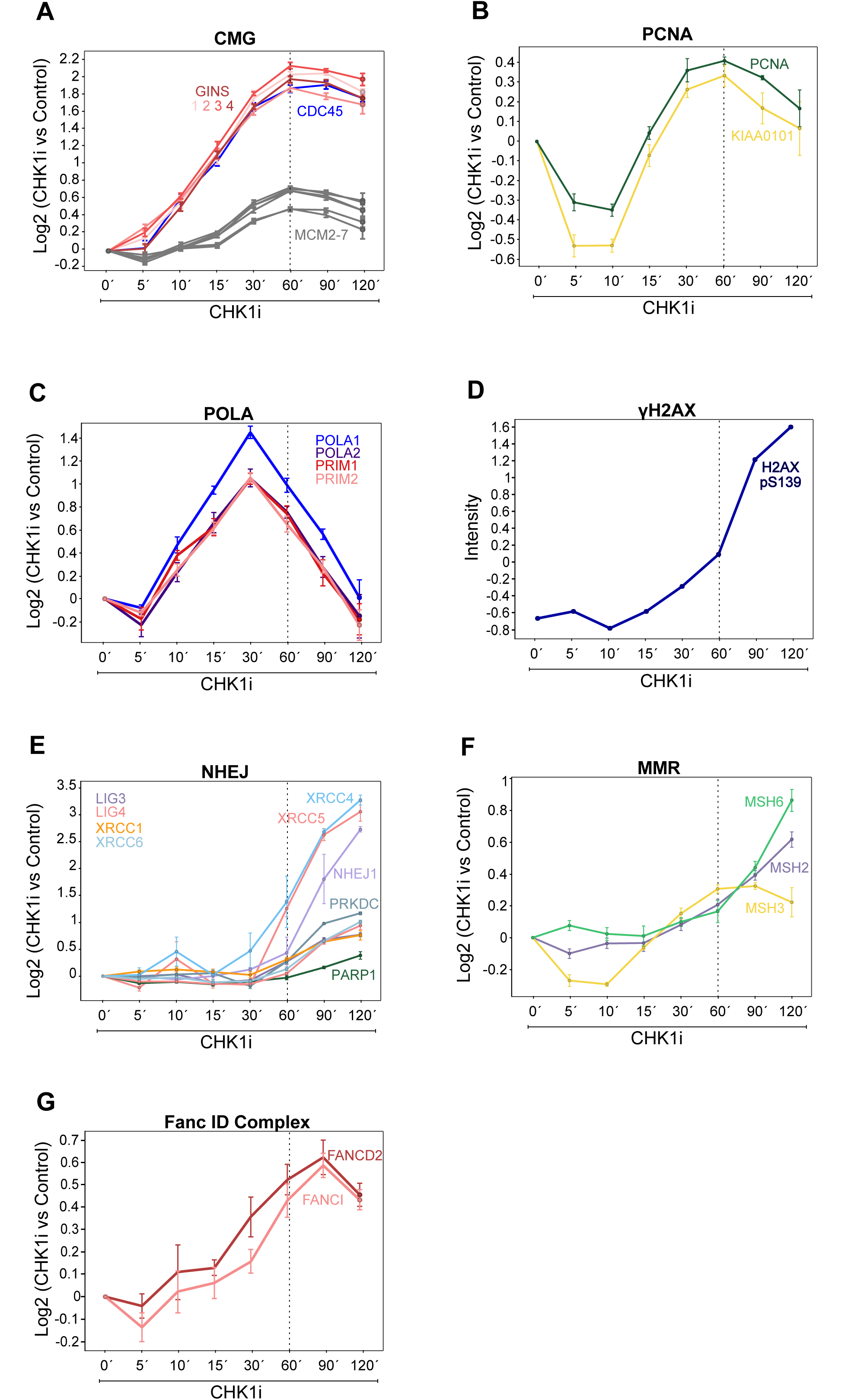
– Time-resolved dynamics of chromatin proteins in CHK1i-treated cells. A-G. Line diagrams showing the logarithmic ratio of chromatin intensities of the indicated factors in CHK1i time course vs control treated cells.

In our analysis, we were also able to identify and quantify 430 phosphorylation sites. To illustrate this, we plotted the absolute intensity of histone H2AX phosphorylated at S139 (γH2AX), a DNA damage marker, in our time-course (Figure 3D and Supplementary Table 2). γH2AX showed only a marginal increase in the earlier time-points and showed a steep rise after 60’ when RC ensued (Figure 3D). This was consistent with the sharp increase in the dynamics of non-homologous end-joining (NHEJ) proteins after the 60’ time-point, indicating the formation of double-strand breaks (DSBs), which are characteristic observed during RC (Figure 3E). However, the levels of homologous recombination (HR) factors did not show a similar increasing trend (Figure S3G). These data indicate that DSBs generated during RC are targeted primarily for repair by the NHEJ pathway. Other DNA repair proteins that are known to be at the vicinity of the replisome, mismatch repair (MMR) and the FANCI-D2 complex, showed loading dynamics that were similar to those of replisome components (Figures 3F-G). Altogether, our method enables probing protein dynamics on the chromatin, as exemplified here by elucidating OF dynamics induced by CHK1i.

### NCAPH2 is a novel protein regulating DNA replication

To further exploit the potential of our dataset, we sought novel factors involved in OF and DNA replication. To do this, we manually selected nearly 50 factors, previously unknown for their role in replication, whose recruitment dynamics mimicked those of known replication proteins and/or were recruited to the chromatin in response to CHK1i (Supplementary Table 3). Next, using QIBC readouts for DNA synthesis (EdU incorporation and chromatin-bound RPA), we performed a targeted siRNA screen on these factors to identify those involved in DNA replication (Figure 4A and Supplementary Table 4). Knockdown of one of these factors, NCAPH2 (Non-SMC Condensin II Complex Subunit H2) showed a reduction in both EdU incorporation and chromatin-bound RPA, similar to what was observed following knockdown of CDC45 (Figure 4A). We were intrigued by this candidate because its chromatin dynamics closely resembled that of PCNA and the PCNA associated factor KIAA0101 (Figure 4B). To validate NCAPH2 as a novel factor involved in DNA replication, we separately transfected cells with four different siRNAs against this protein. In all cases, these cells showed a reduction in DNA synthesis as measured by EdU incorporation (Figure 4C). We repeated this experiment and measured chromatin bound CDC45 as a readout for OF, and found that three out of four siRNAs also resulted in a reduction of CDC45 loading during S phase (Figure 4D). Additionally, dormant OF, induced by treating cells with CHK1i, was also negatively affected in cells transfected with these siRNAs (Figure 4E). To directly assess the importance of NCAPH2 for initiation of replication, we fused a mAID degron tag (mini Auxin-Inducible Degron) onto the C-terminus of NCAPH2 in the near diploid cell line HCT116 TET-TIR1^35^. The cloning was designed to preserve the position and sequence of the NCAPH2 3’ UTR, since it likely has important regulatory functions (Figure S4A, see methods for entire strategy). Insertion of the mAID sequence and blasticidin and hygromycin resistance cassettes within each targeted NCAPH2 allele were confirmed by PCR (Figures S4B-E). The inducible depletion of NCAPH2 upon addition of the synthetic auxin indole-3-acetic acid (IAA) was then confirmed by western blotting (Figure S4F). The fusion protein NCAPH2-mAID localised to the central axis of isolated native mitotic chromosomes (Figure S4G), suggesting that protein function was not compromised^36^. The insertions did not affect the cell cycle distribution compared to parental cells (Figure S4H-I). We used QIBC to directly assess DNA synthesis in S phase of NCAPH2-mAID cells, and we observed a ∼25% decrease in overall DNA synthesis (Figure 4F), as early as 30 minutes after addition of auxin. Correspondingly, CB-RPA levels were decreased in S phase of the depleted cels, confirming a role for NCAPH2 protein in DNA replication (Figure 4G). We also observed a reduction in CB-RPA when origin firing was induced with CHK1i (Figure 4H). Overall, our data strongly suggest a role for NCAPH2 in regulating DNA replication.

**Figure 4.**
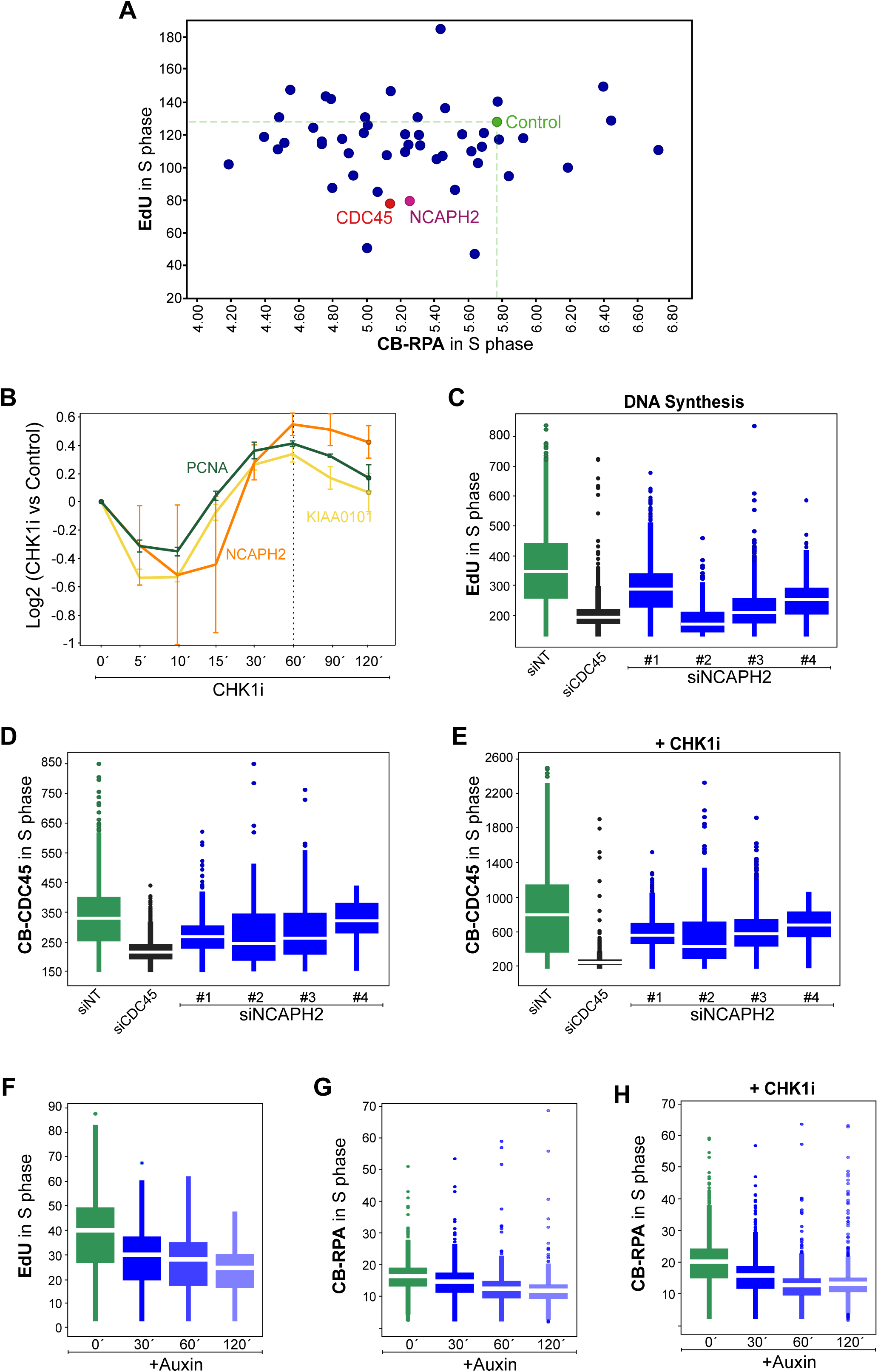
– NCAPH2 is a novel protein regulating DNA replication. A. Targeted siRNA screen on factors selected from the MS dataset (Supplementary Table 3). Average of Mean intensities of EdU and RPA in S phase cells (EdU positive gate) are plotted. Control siRNA is shown in green, CDC45 in red, NCAPH2 in purple and the others in blue. B. Line diagrams showing the dynamics of PCNA, KIAA0101 (PCLAF) and NCAPH2 in our CHK1i time course. C. Cells were transfected with indicated siRNAs for 72h and then stained for EdU. siCDC45 is shown as a positive control. Mean intensities are plotted. White lines indicate median values and outliers are represented as small circles. D. U2OS-eCDC45 cells were transfected with indicated siRNAs for 72h and then stained for GFP and further QIBC analysis was performed. Mean intensities are plotted. White lines indicate median values and outliers are represented as small circles. E. U2OS-eCDC45 cells were transfected with indicated siRNAs for 72h and treated with CHK1i during the last 30 minutes before being stained for GFP and further QIBC analysis was performed. Mean intensities are plotted. White lines indicate median values and outliers are represented as small circles. F. HCT116-Tet-Tir1 cells with NCAPH2 endogenously tagged with mAID (NCAPH2-mAID) were treated with 500µM indole-acetic acid (IAA, auxin) for the indicated times and then stained for EdU and CB-RPA. Mean intensities are plotted. White lines indicate median values and outliers are represented as small circles. G. HCT116-Tet-Tir1 cells with NCAPH2 endogenously tagged with mAID (NCAPH2-mAID) were treated with 500µM indole-acetic acid (IAA, auxin) for the indicated times. During IAA treatment, cells were also treated with CHK1i for 30 minutes. Finally, cells were stained for EdU and CB-RPA. Mean intensities are plotted. White lines indicate median values and outliers are represented as small circles.

## Discussion

In this manuscript, we describe a method to analyse the chromatin-bound proteome of S-phase cells in response to replication origin firing. We provide time-resolved loading dynamics for 3,269 proteins, including the replisome and origin firing factors. Our method was able to identify all the replisome components, even the ones missing from previous proteomic analyses after iPOND or NCC ^34^. Both iPOND and NCC rely on labelling nascent DNA with nucleotide analogues for up to 30 minutes. Generally, replication forks move at a rate of up to 1 kb/min, meaning that the replisome could be 30 kb away from the end of the nascent strand, making it difficult to capture all the replisome proteins. Therefore, iPOND and NCC require large sample quantities and the isolation of nascent strands will also isolate factors that may have no direct role at replication forks. In contrast, our method is technically straightforward and requires only a small sample (fewer than 100,000 cells per replicate). Moreover, our comparison of the CHK1i-induced chromatin proteome to a baseline S phase chromatin proteome revealed factors enriched on the chromatin after origins had fired, but prior to the start of nascent strand synthesis.

Our dataset is an excellent resource of proteins enriched on the chromatin during OF. Over time, many replisome factors were enriched gradually on the chromatin and then their level plateaued, suggesting that at this point no further activation of replication forks was possible. For instance, the essential factor RPA, whose accumulation of chromatin plateaued after 60 minutes of CHK1i treatment, coincided with the sharp increase in yH2AX, which is reminiscent of replication catastrophe (RC) ^17^. Interestingly, we identified NHEJ factors rather than HR factors being recruited to DNA breaks, possibly induced by RPA-exhaustion ^17^. Some core replisome factors showed unexpected trends. PCNA and PCLAF first decreased on chromatin for 10 minutes and only then started to increase. More interesting was the behaviour of the POLA complex, which rapidly unloaded from the chromatin after 30 minutes of CHK1 inhibition, correlating with the putative induction of RC. It could be relevant to investigate a potential functional link of POLA1 and RPA exhaustion after our recent report describing POLA activity in relation to RC ^37^. We also found the FANCI-FANCD2 (ID) complex directly correlating to the number of active replication forks. The ID complex was previously shown to be recruited to the MCMs during replication stress ^27^. It is therefore tempting to speculate that the ID complex is constantly monitoring newly fired forks. Alternatively, the act of origin firing per se, or loss of CHK1 activity might lead to replication stress requiring the recruitment of the ID complex. Further work is required to deduce the role of these proteins on the chromatin during origin firing.

Analysis of chromatin association dynamics highlighted those factors that travel with the replisome, in contrast to other DNA replication proteins that might only be transiently recruited to the chromatin during origin firing. For instance, the components of the core replicative helicase, CDC45 and the GINS complex showed a gradual enrichment on the chromatin, indicating their continuous requirement at active replication forks. In contrast, TOPBP1 and RECQL4, did not enrich on the chromatin to the same relative extent or with the same kinetics as CDC45 upon induction of OF.

Subunits of known complexes in our dataset followed very similar loading dynamics. This reinforces the quantitative power of our method and suggests that proteins with meaningful trends could harbour roles in DNA replication. Using this approach, we chose NCAPH2 for its similarity in loading dynamics with PCNA/PCLAF. Depletion of NCAPH2 affected global DNA replication levels and CDC45 loading. NCAPH2 is a non-SMC subunit of the condensin II complex, known for its role in mitotic DNA compaction and faithful chromosome segregation in mitosis ^38–40^. The condensin II complex has also been implicated in the maintenance of nuclear architecture ^41^ as well as gene regulation in interphase ^42^. Our data suggests a role for a condensin II subunit in DNA replication. It remains to be tested if this is an independent role for NCAPH2 or mediated by the whole condensin II complex.

Altogether, we employed a total chromatin MS-based proteomics approach, which enabled us to uncover the dynamics of proteins during origin firing in human cells. While we cannot rule out that the dynamic loading or unloading of proteins in CHK1i treated cells could be due to inhibition of CHK1 functions other than regulation of origin firing ^43^, our temporal proteomics data allowed us to focus on origin firing by comparing to the dynamics of known replication proteins. Using this technique, we were able to identify a novel factor required for DNA replication, providing a valuable resource for the community. Furthermore, the method described here will allow researchers to investigate other pathways of DNA metabolism in human cells, as well as the effects of other inhibitors and treatments, in a straightforward manner.

## Figure Legends

**Figure S1.**
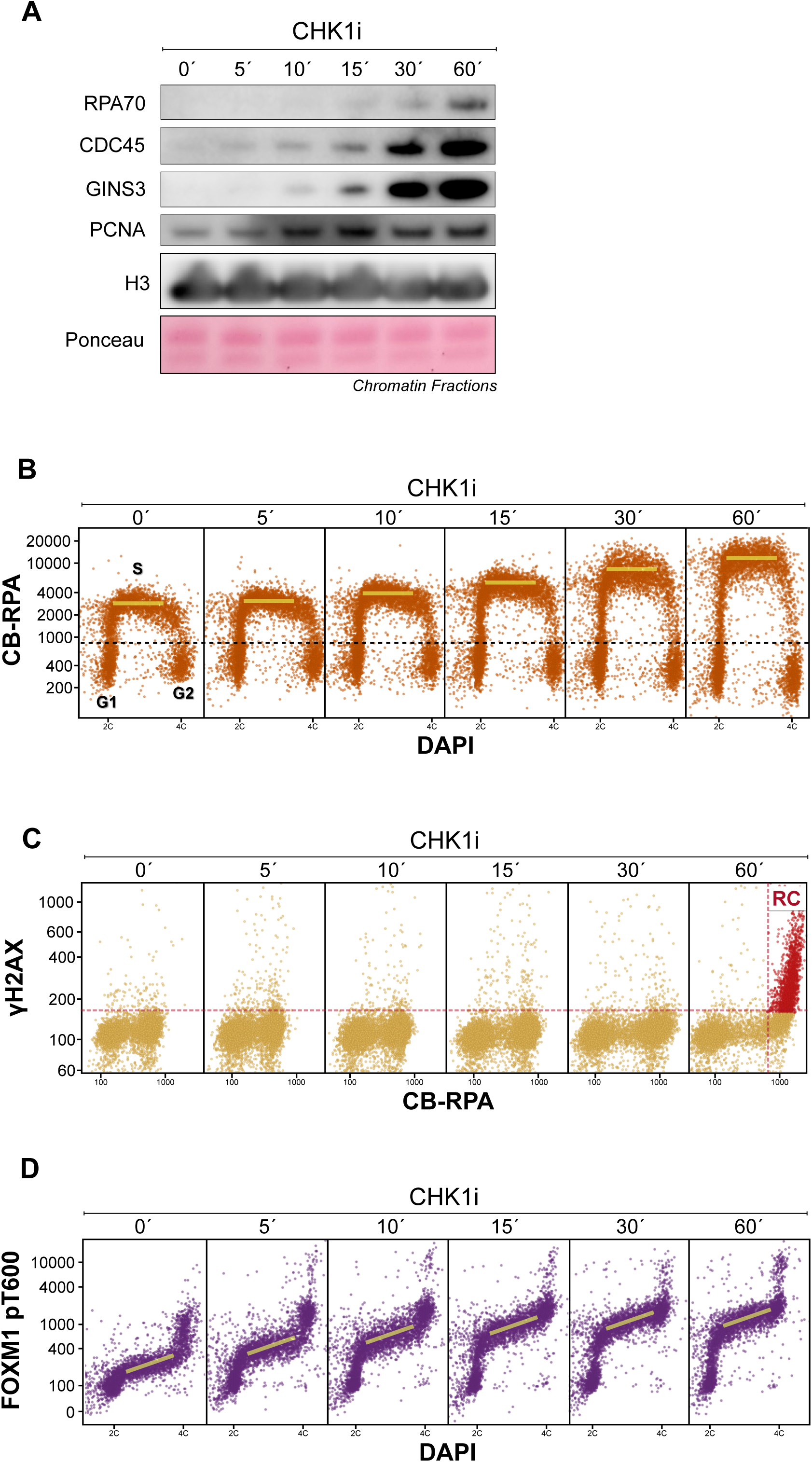
– Related to Figure 1. A. Cells were treated for the indicated times with CHK1i inhibitor and chromatin-enriched fractions were prepared and analysed by western blotting with the indicated antibodies. H3 and Ponceau-S staining were used as a loading control. B. QIBC scatter plots for the experiment in Figure 1B. Cells were gated into G1, S or G2 as indicated. Median CB-RPA intensity in S phase is marked with a yellow line. C. QIBC scatter plots for CB-RPA/γH2AX for the experiment in Figure 1B. Cells in RC (red) were marked as previously described ^17,44^. D. QIBC scatter plots for the experiment in Figure 1E. The trend of FOXM1 pT600 in S phase cells is marked with a yellow line.

**Supplementary Figure 2.**
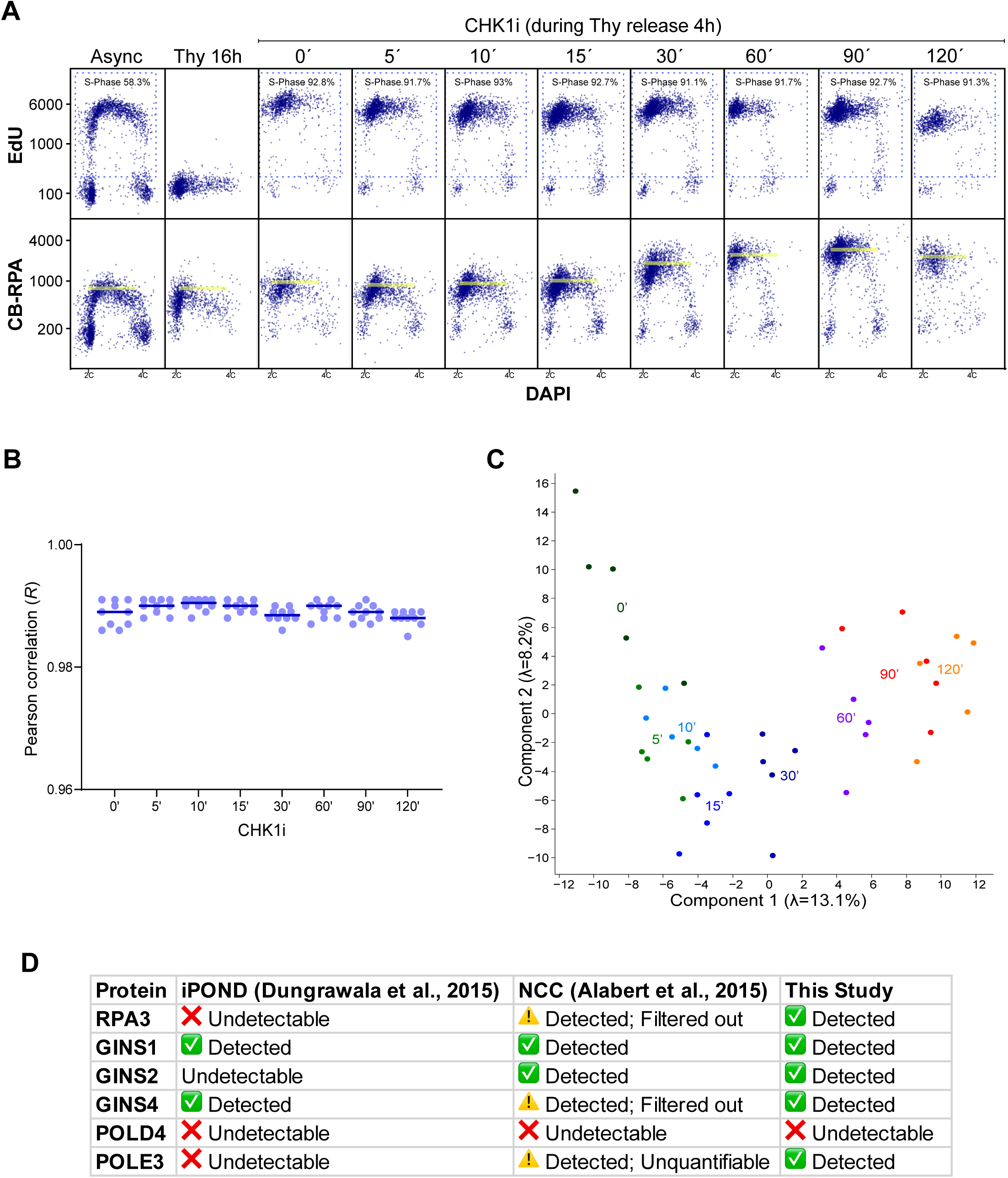
– Related to Figure 2. A. QIBC scatter plots for EdU/DAPI (top panel) and RPA/DAPI (bottom panel) in cells processed as in Figure 2A. Only one replicate is shown. EdU positive cells were gated as S phase cells and quantified. In the bottom panel, the peak of RPA is indicated with a yellow line. B. Comparison of some critical replisome components quantified by recent iPOND and NCC studies with our study. Adapted from Cortez, 2017 ^34^.

**Supplementary Figure 3.**
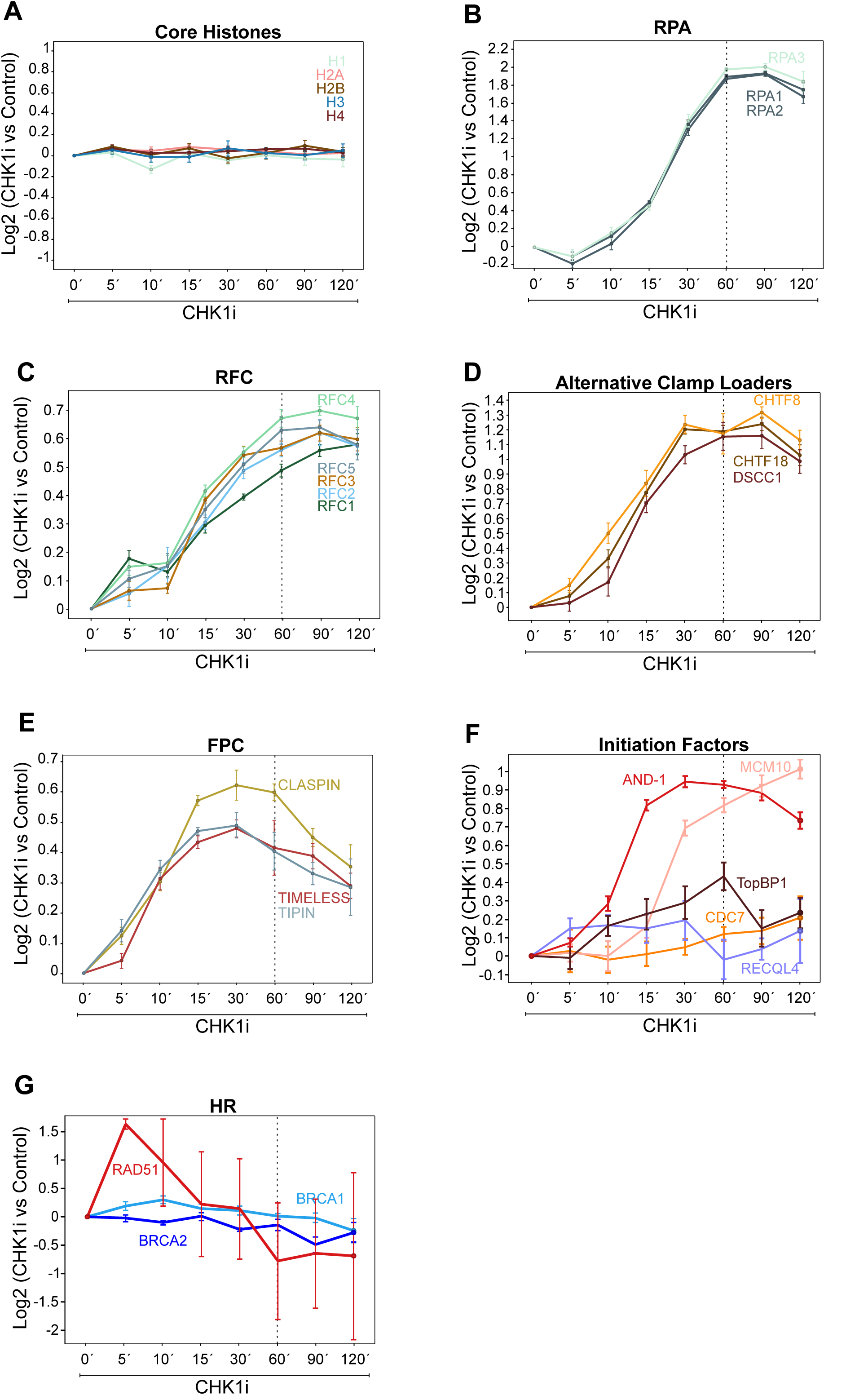
– Related to Figure 3. A-G. Line diagrams showing the logarithmic ratio of chromatin intensities of the indicated factors in CHK1i time course vs control treated cells.

**Supplementary Figure 4.**
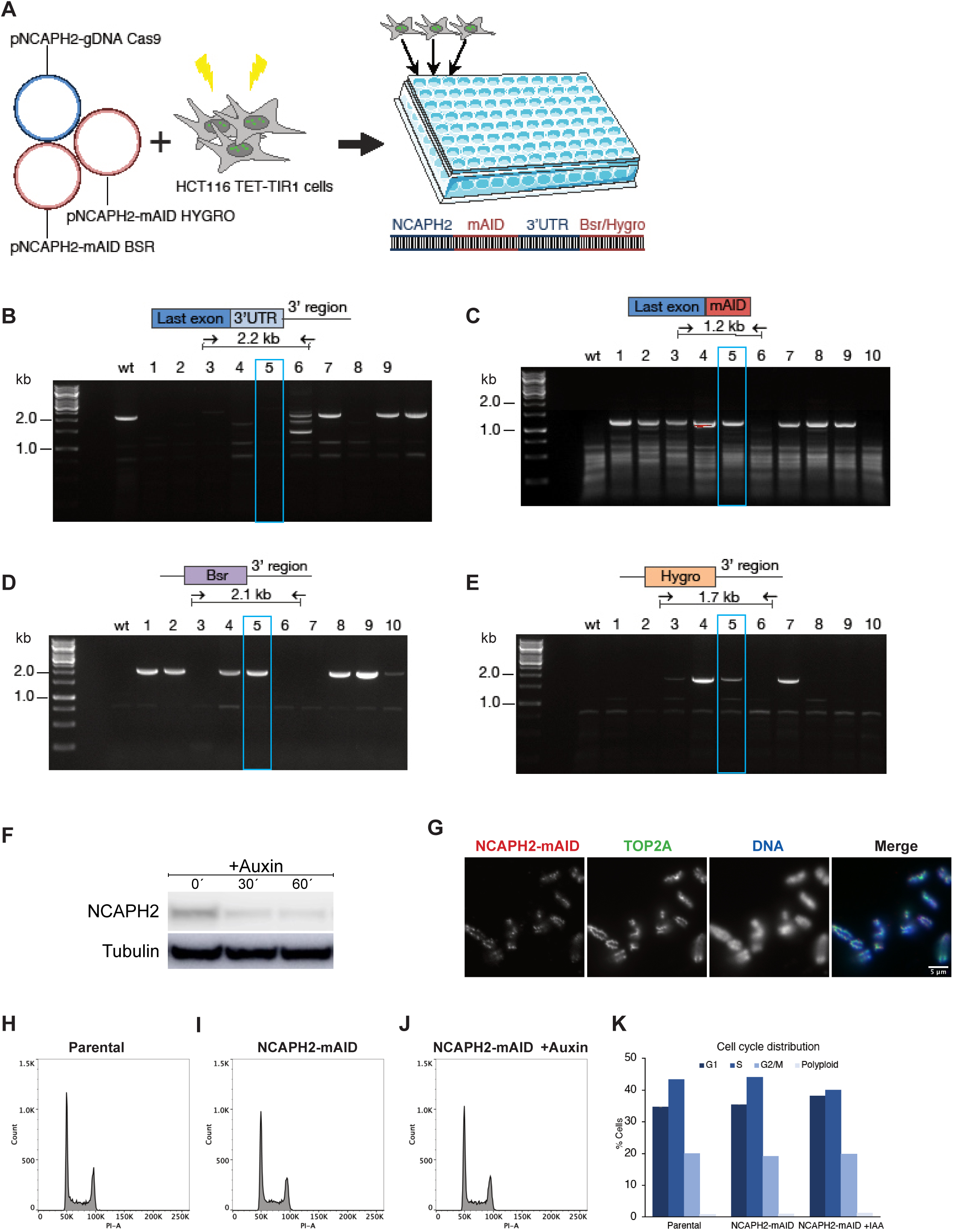
– Related to Figure 4 – Development of an auxin-inducible NCAPH2 degron cell line. A. Diagram of the cloning strategy. Plasmids encoding Cas9 and guide targeting the 3’ UTR of the NCAPH2 gene, and homology directed repair templates for insertion of mAID and Bsr/Hygro cassettes was transfected into HCT116 TET-TIR1 cells. Single clones were selected and the entire 3’UTR of NCAPH2 was preserved following mAID insertion at the C terminal like depicted in the sequence diagram. B-E. Agarose gel images of genomic PCR products from wildtype (wt) and NCAPH2-mAID clone DNA, amplifying the targeted locus (B) or overlaps between the last exon and mAID (C), the Bsr Cassette and 3’UTR (D) and the Hygro cassette and 3’UTR (E). The blue boxes indicate the chosen correct clone (5). F. Western blot of cells treated with Doxycycline (18 hours) and Auxin (indicated times) and stained with NCAPH2 antibody. Tubulin was used as a loading control. G. Native mitotic chromosomes from NCAPH2-mAID cells were immunostained for NCAPH2 (red) and Topo II (green). DNA was stained with DAPI (blue). H. Flow cytometry histograms of the asynchronous cell cycle distribution of the parental TET-TIR1 cells, CAPH2-mAID cells or CAPH2-mAID cells treated with Doxycycline (18 hours) and Auxin (4 hours). I. Quantification of the cell cycle distributions from H.

## Methods

### Cell Culture

All experiments were performed in U2OS cells. For assessing CB-CDC45 via QIBC, U2OS cells with CDC45 endogenously tagged with GFP (U2OS-eCDC45; kind gift from Jiri Lukas) were used. All cells were cultured at 37°C in DMEM, supplemented with 6% foetal bovine serum (FBS) and Penicillin-Streptomycin (10,000 U/mL) all purchased from Thermo Fisher Scientific. The cell lines were not authenticated and were verified to be free of mycoplasma contamination once every few months. CHK1i (AZD7762, Selleckchem) was used at a concentration of 1µM. Thymidine (Sigma-Aldrich) was used at 2mM to synchronise cells at the G1/S boundary. CDC7 inhibitor XL413 (Selleckchem) was used at a concentration of 1 µM. Pan CDK inhibitor R547 (Adooq Bioscience) was also used at a concentration of 1 µM.

### QIBC

QIBC was essentially performed as described before ^17^. Cells in 96-well plates were placed on ice and either pre-extracted (to remove soluble proteins) or not with 0.5% Triton X-100 PBS on ice for 1 min before fixing with 4% paraformaldehyde. Fixed cells were washed with 0.01% Tween-20 PBS and incubated with primary and later secondary antibodies diluted in filtered complete DMEM. Primary antibodies for RPA (1/1000, custom made, detects all RPA subunits); γH2AX (1/5000, JBW301, Merck Millipore); pFOXM1 (1/500, Cell Signaling) and GFP (1/2000, Cromotek) were diluted in filtered complete cell culture medium and incubated at room temperature for 1 hour. For detecting EdU, Click-it reaction was performed after staining with primary antibodies by incubating cells in Click-it buffer (100 mM Tris-HCl pH 8, 2 mM CuSO4, 1 ng Alexa Fluor 647 Azide (Life Technologies), and 100 mM ascorbic acid for 30 min at room temperature (RT). Secondary antibodies (Alexa Fluor Plus) were also diluted in filtered completed medium and incubated for 1 hour at room temperature. After each round of antibody incubation, cells were washed thrice with 0.01% Tween-20 PBS. Images for QIBC were automatically obtained using a motorised Olympus IX-83 wide-field ScanR microscope and the accompanying acquisition software. The microscope was equipped with filter cubes compatible with DAPI, FITC, Cy3, and Cy5 fluorescent dyes, aSpectra X-LIGHT engine Illumination system with 6 colour LEDs and emission filters, and a Hamamatsu Camera Orca Flash4.0 V2. An Olympus Universal Plan Super Apo 10x Objective was used for all QIBC data. Acquired images were processed with the ScanR image analysis program. TIBCO Spotfire® software was used to perform cytometry and analyse nuclear pixel intensities in total for DAPI (Arbitrary units: A.U.) and mean (total intensity divided by nuclear area; also A.U.) for all other parameters (unless otherwise specified) and visualised as scatter plots.

### Western Blotting

Whole cell extracts (WCE) were obtained by lysis in RIPA buffer (50 mM Tris-HCL pH 8.0, 150 mM NaCl, 1.0% IGEPAL CA-630, 0.1% SDS, and 0.1% Na-deoxycholic acid) supplemented with protease and phosphatase inhibitors (Roche) and Benzonase® Nuclease (Sigma-Aldrich) for 30 min. WCE, soluble and chromatin fractions were analysed by SDS–PAGE after boiling samples in reducing buffer (DTT; Sigma-Aldrich) as per standard procedures. For immunoblotting, membranes were blocked in a blocking buffer (0.05% Tris Buffered Saline – Tween20 containing 5% skimmed milk powder) before incubating with primary antibodies overnight at 4°C in the same blocking buffer. Secondary peroxidase-coupled antibodies were also incubated in blocking buffer at RT for 1h. ECL-based chemiluminescence was detected using an Amersham Imager 600. Primary antibodies were used at the indicated dilutions: CDC45 (1/1000, Cell Signaling); GINS3 (1/3000, Bethyl Laboratories); Histone H3 (1/3000, Abcam); PCNA (1/500, Santa Cruz); TopBP1 (1/1000, Bethyl Laboratories); RPA70 (1/1000, Abcam); RECQL4 (1/1000, Novus Biologicals); TRESLIN (Custom antibody, kind gift from Dominic Boos); Vinculin (1/10000, Sigma); γH2AX (1/5000, JBW301, Merck Millipore); NCAPH2 (1:1000, sc-393333 Santa Cruz), α-Tubulin (1:10000, ab18251 Abcam).

### Preparation of chromatin-enriched fractions

Cells were washed with ice-cold PBS twice and collected by scraping. Soluble fraction was extracted by lysing the cells with an extraction buffer (0.5% Triton X-100 in PBS) for 20 minutes on ice. Supernatant containing the soluble pool of proteins was collected. The remaining pellet (chromatin-enriched fraction) was washed again with the extraction buffer for 5 minutes before finally washing thrice with cold PBS. For western blotting, pellets were lysed in RIPA buffer (50mM Tris pH8.0, 150mM NaCl, 1% Igepal CA-630, 0.1% SDS, 0.1% Sodium-deoxycholic acid, 250U/mL Benzonase nuclease). For MS, pellets were lysed in 6M Guanidine-HCl to generate chromatin samples.

### Chromatin Sample Preparation for MS

Chromatin samples were reduced and alkylated by addition of tris(2-carboxyethyl)phosphine and chloroacetamide to final concentrations of 5 mM, after which they were digested using endoproteinase Lys-C (1:200 w/w; Wako Chemicals), for 4 hours at room temperature. Samples were diluted 4-fold using 50 mM ammonium bicarbonate, after which they were further digested using sequencing grade Trypsin (1:100 w/w; Sigma), overnight at room temperature. Samples were clarified of minor precipitation by passing them through 0.45 μm spin filters. Tryptic peptides were purified and fractionated on-StageTip at high-pH essentially as described previously ^45^. C18 StageTips were prepared in-house, by layering four plugs of C18 material (Sigma-Aldrich, Empore™ SPE Disks, C18, 47 mm) per StageTip. Activation of StageTips was performed with 100 μL 100% methanol, followed by equilibration using 100 μL 80% acetonitrile (ACN) in 200 mM ammonium hydroxide, and two washes with 100 μL 50 mM ammonium hydroxide. Samples were basified by addition of 1/10th volume of 200 mM ammonium hydroxide, after which they were loaded on StageTips. Subsequently, StageTips were washed twice using 100 μL of 50 mM ammonium hydroxide, after which peptides were eluted as four fractions (F1-4) using 3%, 7%, 13%, and 25% ACN in 50 mM ammonium hydroxide. All fractions were dried to completion using a SpeedVac at 60 °C. Dried peptides were dissolved in 20 μL 0.1% formic acid, and stored at −20°C, until analysis of 5 μL from each sample using mass spectrometry.

### MS Data Acquisition

All samples were analysed on an EASY-nLC 1200 system (Thermo) coupled to an Orbitrap Exploris™ 480 mass spectrometer (Thermo). Separation of peptides was achieved using 15-cm columns (75 µm internal diameter) packed in-house with ReproSil-Pur 120 C18-AQ 1.9 µm beads (Dr. Maisch). Elution of peptides from the column was achieved using a gradient ranging from buffer A (0.1% formic acid) to buffer B (80% acetonitrile in 0.1% formic acid), at a flow rate of 250 nL/min. Gradient length was 80 min per sample, including ramp-up and wash-out, with an analytical gradient of 57 min ranging in buffer B from 5-32% for F1, 5-36% for F2, 6-38% for F3, 7-40% for F4. The column was heated to 40°C using a column oven, and ionization was achieved using a NanoSpray Flex™ NG ion source (Thermo). Spray voltage set at 2 kV, ion transfer tube temperature to 275°C, and RF funnel level to 40%. Full scan range was set to 300-1,300 m/z, MS1 resolution to 120,000, MS1 AGC target to “200” (2,000,000 charges), and MS1 maximum injection time to “Auto”. Data was acquired in data-dependent mode, and the top 18 most abundant precursors were acquired each duty cycle. Precursors with charges 2-6 were selected for fragmentation using an isolation width of 1.3 m/z and were fragmented using higher-energy collision dissociation (HCD) with normalised collision energy of 25. Monoisotopic Precursor Selection (MIPS) was enabled in “Peptide” mode, without relaxing restrictions in case too few precursors were observed. Precursors were prevented from being repeatedly sequenced by setting expected peak width to 40 s, and setting dynamic exclusion duration to 80 s, with an exclusion mass tolerance of 15 ppm, exclusion of isotopes, and exclusion of alternate charge states for the same precursor. MS/MS resolution was set to 15,000, MS/MS AGC target to “200” (200,000 charges), MS/MS intensity threshold to 430,000, MS/MS maximum injection time to “Auto”, and first mass for the MS/MS scan range was set to 100 m/z.

### MS Data Analysis

All MS RAW data were analysed using the freely available MaxQuant software ^46,47^ v.1.5.3.30. Default MaxQuant settings were used, with exceptions specified below. For generation of theoretical spectral libraries, the human FASTA database was downloaded from UniProt on the 24th of May 2019. In-silico digestion of proteins to generate theoretical peptides was performed with trypsin, allowing up to 6 missed cleavages. Allowed variable modifications were oxidation of methionine (default) and protein N-terminal acetylation (default), with maximum variable modifications per peptide reduced to 3. First search mass tolerance was reduced to 10 ppm, and maximum precursor charge to consider was reduced to 6. Second peptide search was enabled. Matching between runs was enabled, with an alignment window of 20 min and a match time window of 1 min, and with matching across different fractions disallowed. Label-free quantification (LFQ) was enabled ^48^, with “Fast LFQ” disabled. iBAQ quantification was enabled. Stringent MaxQuant 1% FDR data filtering at the PSM– and protein-levels was applied (default).

### MS Data Annotation and Quantification

Quantification of the MaxQuant output files (“proteinGroups.txt”) was performed using Perseus software ^49^. Experiments were performed in biological quadruplicate, with a single replicate prepared as one batch, and three further replicates as a separate batch. As a precaution against batch-to-batch variance, an equimolar mixture was created from the three same-batch replicates just prior to MS analysis and measured as a fifth replicate. For quantification purposes, all protein LFQ intensity values were log2 transformed. Proteins were first filtered for a presence in 4 out of 5 replicates in every condition, after which any missing values were imputed using the ImputeLCMD R package as integrated in Perseus (https://cran.r-project.org/web/packages/imputeLCMD/imputeLCMD.pdf) and using the Singular Value Decomposition (SVD) algorithm with a minimum of two principal components. Proteins that were not present in 4 out of 5 replicates in every condition were retained in a separate list, which was then filtered for proteins to be present in 5 out of 5 replicates in at least one experimental condition. In this case, missing values were imputed from a normal distribution, below the global experimental detection limit at a downshift of 1.8 and a randomised width of 0.15 (in log2 space). Following this, the two lists of proteins were merged, and any proteins not matching either of the filters (n=4/5 in all conditions, or n=5/5 in any condition) were considered unquantifiable and omitted. Subsequently, we compensated the data for the presence of batch effects, as replicate #1 was prepared separately from replicates #2-3-4, and furthermore replicate #5 was measured on a different instrument due to technical complications. Batch effort removal was performed using the “Remove batch effect” R package as integrated into Perseus, using the ComBat algorithm ^50^, and with three experimental batches defined as indicated above. Unsupervised hierarchical clustering and expression profile analyses were performed after z-scoring the data. Statistical significance of differences was tested using two-tailed Student’s t testing, with permutation-based FDR-control applied at an s0 value of 0.1.

The mass spectrometry proteomics data have been deposited to the ProteomeXchange Consortium via the PRIDE ^51^ partner repository with the dataset identifier PXD047108.

### Generation of NCAPH2-mAID cell line

The guide DNA sequence 5’-GCCTCGGTGCTCCCCACTCA (PAM: GGG, NCAPH2 gDNA 1 forward and reverse) targeting the 3’UTR of NCAPH2 3’UTR was ligated into the pSpCas9(BB)– 2A-GFP (pX458) plasmid ^52^. For template plasmids the following fragments were joined by Gibson assembly (NEBuilder HiFi DNA assembly®, NEB): pUC19*EcorI (NEB), 5’ homology arm-Linker-mAID-3’UTR, BSR/Hygro cassettes, 3’ homology arm. The 3’ homology arm was amplified by PCR (Q5 HiFi polymerase®, NEB), from HCT116 cell genomic DNA using the primers NCAPH2 3’ homology arm forward and reverse, and gel-purified. The Blasticidin or Hygromycin cassettes were cut and gel-purified from either pMK288 or pMK287 plasmids ^35^ (kind gifts from Prof. M. Kanemaki, RIKEN) using MfeI and BamHI restriction enzymes. The 5’ homology arm-Linker-mAID-3’UTR fragment was synthesized using Genestrings® (Thermo Fisher Scientific). The Cas9-NCAPH2-gDNA plasmid and the two template plasmids (BSR or Hygro) were transfected into HCT116 TET-TIR1 cells ^35^ using Neon Electroporation System® (Thermo Fisher Scientific) according to manufacturer instructions. The day after transfection cells were treated with 125μg/mL hygromycin and 7.5μg/mL blasticidin for more than 12 days before colony selection. Correct clones were confirmed by PCR (Figure S4B-E), sequencing (Macrogen), immunoblotting (Figure S4F) and immunofluorescent microscopy on chromosome spreads (Figure S4G) like described before ^53^. All DNA sequences used for the generation of the NCAPH2-mAID cell line are deposited in Table 1.

**Table 1.**
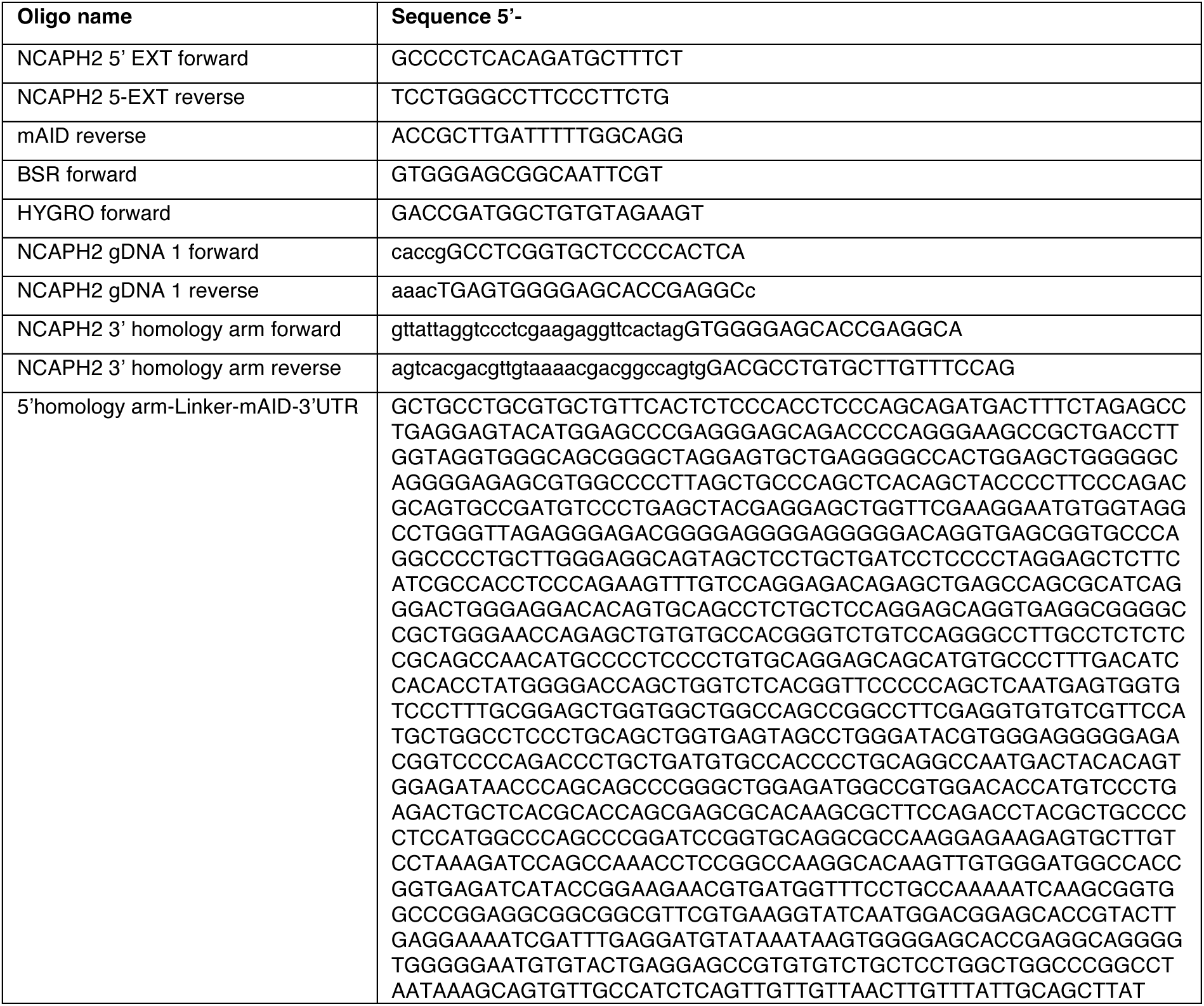
A table with primers and oligos used in the study.

### Native chromosome immunofluorescence

Native chromosomes were isolated like described previously ^54^. All steps were done at room temperature. Chromosome isolate was diluted 1:3 in chromosome buffer (15mM Tris-HCl pH 7.4, 0.5mM EDTA-K, 75mM KCl, 1mM spermidine, 0.4mM spermine). and then spotted on poly-L-lysine coated glass 1.5# coverslips for 10 min. Coverslips were washed once gently in chromosome buffer with 3% BSA and then incubated for 1 hour. Coverslips were then incubated with primary antibodies for two hours at room temperature, washed three times in chromosome buffer with 3% BSA, incubated 1 hour with secondary antibodies in chromosome buffer with 3% BSA, washed twice with chromosome buffer, incubated 10 min with 100ng/mL DAPI in chromosome buffer, then washed three times in chromosome buffer. Coverslips were mounted on superfrosted slides in 5X chromosome buffer diluted to 1X in Vectashield® antifade mounting medium. Primary antibodies were TOP2A 1:400 (sc-5347, Santa Cruz), NCAPH2 1:100 (sc-393333, Santa Cruz) with the corresponding secondary antibodies 1:1000 alexa fluor® 647 donkey anti-goat (A21447, Invitrogen) 1:1000 alexa fluor® 568 donkey anti-mouse (A10037, Invitrogen). Slides were images using a 60X objective with 1.4 NA mounted on a BX63 microscope (Olympus).

## Acknowledgements

We would like to thank Dr Jiri Lukas and his lab for the U2OS endogenously tagged-CDC45 cells. We thank all authors and members of the Toledo lab for critical reading, revising and providing feedback on the manuscript. Work at the Center for Chromosome Stability was supported by the European Research Council, European Union (ERC-StG-679754), and the Danish National Research Foundation, Denmark (DNRF115). Work in the MLN lab was supported by the Independent Research Fund Denmark (0135-00096B and 8020-00220B), the European Union’s Horizon 2020 research and innovation program under grant agreement EPIC-XS-823839, and the Danish Cancer Society (R146-A9159-16-S2).

## Declaration of interests

The authors declare no competing interests.

## Notes

### Competing Interest Statement

The authors have declared no competing interest.

## References

1. Yeeles, J.T.P., Deegan, T.D., Janska, A., Early, A., and Diffley, J.F.X. (2015). Regulated eukaryotic DNA replication origin firing with purified proteins. Nature 519, 431–435. 10.1038/nature14285.

2. Baris, Y., Taylor, M.R.G., Aria, V., and Yeeles, J.T.P. (2022). Fast and efficient DNA replication with purified human proteins. Nature 606, 204–210. 10.1038/s41586-022-04759-1.

3. Remus, D., Beuron, F., Tolun, G., Griffith, J.D., Morris, E.P., and Diffley, J.F.X. (2009). Concerted loading of Mcm2-7 double hexamers around DNA during DNA replication origin licensing. Cell 139, 719–730. 10.1016/j.cell.2009.10.015.

4. Padayachy, L., Ntallis, S.G., and Halazonetis, T.D. (2024). RECQL4 is not critical for firing of human DNA replication origins. Sci. Rep. 14. 10.1038/s41598-024-58404-0.

5. Boos, D., Sanchez-Pulido, L., Rappas, M., Pearl, L.H., Oliver, A.W., Ponting, C.P., and Diffley, J.F.X. (2011). Regulation of DNA replication through Sld3-Dpb11 interaction is conserved from yeast to humans. Curr. Biol. 21, 1152–1157. 10.1016/j.cub.2011.05.057.

6. Boos, D., Yekezare, M., and Diffley, J.F.X. (2013). Identification of a heteromeric complex that promotes DNA replication origin firing in human cells. Science 340, 981–984. 10.1126/science.1237448.

7. Im, J.-S., Ki, S.-H., Farina, A., Jung, D.-S., Hurwitz, J., and Lee, J.-K. (2009). Assembly of the Cdc45-Mcm2-7-GINS complex in human cells requires the Ctf4/And-1, RecQL4, and Mcm10 proteins. Proc. Natl. Acad. Sci. U. S. A. 106, 15628–15632. 10.1073/pnas.0908039106.

8. Kumagai, A., Shevchenko, A., Shevchenko, A., and Dunphy, W.G. (2010). Treslin collaborates with TopBP1 in triggering the initiation of DNA replication. Cell 140, 349–359. 10.1016/j.cell.2009.12.049.

9. Sangrithi, M.N., Bernal, J.A., Madine, M., Philpott, A., Lee, J., Dunphy, W.G., and Venkitaraman, A.R. (2005). Initiation of DNA replication requires the RECQL4 protein mutated in Rothmund-Thomson syndrome. Cell 121, 887–898. 10.1016/j.cell.2005.05.015.

10. Douglas, M.E., Ali, F.A., Costa, A., and Diffley, J.F.X. (2018). The mechanism of eukaryotic CMG helicase activation. Nature 555, 265–268. 10.1038/nature25787.

11. Tanaka, S., Umemori, T., Hirai, K., Muramatsu, S., Kamimura, Y., and Araki, H. (2007). CDK-dependent phosphorylation of Sld2 and Sld3 initiates DNA replication in budding yeast. Nature 445, 328–332. 10.1038/nature05465.

12. Zegerman, P., and Diffley, J.F.X. (2007). Phosphorylation of Sld2 and Sld3 by cyclin-dependent kinases promotes DNA replication in budding yeast. Nature 445, 281–285. 10.1038/nature05432.

13. Burgers, P.M.J., and Kunkel, T.A. (2017). Eukaryotic DNA Replication Fork. Annu. Rev. Biochem. 86, 417–438. 10.1146/annurev-biochem-061516-044709.

14. Choe, K.N., and Moldovan, G.-L. (2017). Forging Ahead through Darkness: PCNA, Still the Principal Conductor at the Replication Fork. Mol. Cell 65, 380–392. 10.1016/j.molcel.2016.12.020.

15. Johnson, R.E., Klassen, R., Prakash, L., and Prakash, S. (2015). A Major Role of DNA Polymerase δ in Replication of Both the Leading and Lagging DNA Strands. Mol. Cell 59, 163–175. 10.1016/j.molcel.2015.05.038.

16. Terret, M.-E., Sherwood, R., Rahman, S., Qin, J., and Jallepalli, P.V. (2009). Cohesin acetylation speeds the replication fork. Nature 462, 231–234. 10.1038/nature08550.

17. Toledo, L.I., Altmeyer, M., Rask, M.-B., Lukas, C., Larsen, D.H., Povlsen, L.K., Bekker-Jensen, S., Mailand, N., Bartek, J., and Lukas, J. (2013). ATR prohibits replication catastrophe by preventing global exhaustion of RPA. Cell 155, 1088–1103. 10.1016/j.cell.2013.10.043.

18. Zhu, W., Ukomadu, C., Jha, S., Senga, T., Dhar, S.K., Wohlschlegel, J.A., Nutt, L.K., Kornbluth, S., and Dutta, A. (2007). Mcm10 and And-1/CTF4 recruit DNA polymerase alpha to chromatin for initiation of DNA replication. Genes Dev. 21, 2288–2299. 10.1101/gad.1585607.

19. Somyajit, K., Gupta, R., Sedlackova, H., Neelsen, K.J., Ochs, F., Rask, M.-B., Choudhary, A., and Lukas, J. (2017). Redox-sensitive alteration of replisome architecture safeguards genome integrity. Science 358, 797–802. 10.1126/science.aao3172.

20. Bermudez, V.P., Maniwa, Y., Tappin, I., Ozato, K., Yokomori, K., and Hurwitz, J. (2003). The alternative Ctf18-Dcc1-Ctf8-replication factor C complex required for sister chromatid cohesion loads proliferating cell nuclear antigen onto DNA. Proc. Natl. Acad. Sci. U. S. A. 100, 10237–10242. 10.1073/pnas.1434308100.

21. Moiseeva, T., Hood, B., Schamus, S., O’Connor, M.J., Conrads, T.P., and Bakkenist, C.J. (2017). ATR kinase inhibition induces unscheduled origin firing through a Cdc7-dependent association between GINS and And-1. Nat. Commun. 8, 1392. 10.1038/s41467-017-01401-x.

22. Moiseeva, T.N., and Bakkenist, C.J. (2018). Regulation of the initiation of DNA replication in human cells. DNA Repair 72, 99–106. 10.1016/j.dnarep.2018.09.003.

23. Sirbu, B.M., Couch, F.B., Feigerle, J.T., Bhaskara, S., Hiebert, S.W., and Cortez, D. (2011). Analysis of protein dynamics at active, stalled, and collapsed replication forks. Genes Dev. 25, 1320–1327. 10.1101/gad.2053211.

24. Alabert, C., Bukowski-Wills, J.-C., Lee, S.-B., Kustatscher, G., Nakamura, K., de Lima Alves, F., Menard, P., Mejlvang, J., Rappsilber, J., and Groth, A. (2014). Nascent chromatin capture proteomics determines chromatin dynamics during DNA replication and identifies unknown fork components. Nat. Cell Biol. 16, 281–293. 10.1038/ncb2918.

25. Dungrawala, H., Rose, K.L., Bhat, K.P., Mohni, K.N., Glick, G.G., Couch, F.B., and Cortez, B. (2015). The Replication Checkpoint Prevents Two Types of Fork Collapse without Regulating Replisome Stability. Mol. Cell 59, 998–1010. 10.1016/j.molcel.2015.07.030.

26. Lopez-Contreras, A.J., Ruppen, I., Nieto-Soler, M., Murga, M., Rodriguez-Acebes, S., Remeseiro, S., Rodrigo-Perez, S., Rojas, A.M., Mendez, J., Muñoz, J., et al. (2013). A proteomic characterization of factors enriched at nascent DNA molecules. Cell Rep. 3, 1105– 1116. 10.1016/j.celrep.2013.03.009.

27. Lossaint, G., Larroque, M., Ribeyre, C., Bec, N., Larroque, C., Décaillet, C., Gari, K., and Constantinou, A. (2013). FANCD2 binds MCM proteins and controls replisome function upon activation of s phase checkpoint signaling. Mol. Cell 51, 678–690. 10.1016/j.molcel.2013.07.023.

28. Sirbu, B.M., McDonald, W.H., Dungrawala, H., Badu-Nkansah, A., Kavanaugh, G.M., Chen, Y., Tabb, D.L., and Cortez, D. (2013). Identification of Proteins at Active, Stalled, and Collapsed Replication Forks Using Isolation of Proteins on Nascent DNA (iPOND) Coupled with Mass Spectrometry*. J. Biol. Chem. 288, 31458–31467. 10.1074/jbc.M113.511337.

29. Zabludoff, S.D., Deng, C., Grondine, M.R., Sheehy, A.M., Ashwell, S., Caleb, B.L., Green, S., Haye, H.R., Horn, C.L., Janetka, J.W., et al. (2008). AZD7762, a novel checkpoint kinase inhibitor, drives checkpoint abrogation and potentiates DNA-targeted therapies. Mol. Cancer Ther. 7, 2955–2966. 10.1158/1535-7163.MCT-08-0492.

30. Morgan, M.A., Parsels, L.A., Zhao, L., Parsels, J.D., Davis, M.A., Hassan, M.C., Arumugarajah, S., Hylander-Gans, L., Morosini, D., Simeone, D.M., et al. (2010). Mechanism of radiosensitization by the Chk1/2 inhibitor AZD7762 involves abrogation of the G2 checkpoint and inhibition of homologous recombinational DNA repair. Cancer Res. 70, 4972– 4981. 10.1158/0008-5472.CAN-09-3573.

31. Bartek, J., and Lukas, J. (2003). Chk1 and Chk2 kinases in checkpoint control and cancer. Cancer Cell 3, 421–429. 10.1016/S1535-6108(03)00110-7.

32. Daigh, L.H., Liu, C., Chung, M., Cimprich, K.A., and Meyer, T. (2018). Stochastic Endogenous Replication Stress Causes ATR-Triggered Fluctuations in CDK2 Activity that Dynamically Adjust Global DNA Synthesis Rates. Cell Systems 7, 17–27.e3. 10.1016/j.cels.2018.05.011.

33. Wessel, S.R., Mohni, K.N., Luzwick, J.W., Dungrawala, H., and Cortez, D. (2019). Functional Analysis of the Replication Fork Proteome Identifies BET Proteins as PCNA Regulators. Cell Rep. 28, 3497–3509.e4. 10.1016/j.celrep.2019.08.051.

34. Cortez, D. (2017). Chapter Two – Proteomic Analyses of the Eukaryotic Replication Machinery. In Methods in Enzymology, B. F. Eichman, ed. (Academic Press), pp. 33–53. 10.1016/bs.mie.2017.03.002.

35. Natsume, T., Kiyomitsu, T., Saga, Y., and Kanemaki, M.T. (2016). Rapid Protein Depletion in Human Cells by Auxin-Inducible Degron Tagging with Short Homology Donors. Cell Rep. 15, 210–218. 10.1016/j.celrep.2016.03.001.

36. Ono, T., Losada, A., Hirano, M., Myers, M.P., Neuwald, A.F., and Hirano, T. (2003). Differential contributions of condensin I and condensin II to mitotic chromosome architecture in vertebrate cells. Cell 115, 109–121. 10.1016/s0092-8674(03)00724-4.

37. Ercilla, A., Benada, J., Amitash, S., Zonderland, G., Baldi, G., Somyajit, K., Ochs, F., Costanzo, V., Lukas, J., and Toledo, L. (2020). Physiological Tolerance to ssDNA Enables Strand Uncoupling during DNA Replication. Cell Rep. 30, 2416–2429.e7. 10.1016/j.celrep.2020.01.067.

38. Cutts, E.E., and Vannini, A. (2020). Condensin complexes: understanding loop extrusion one conformational change at a time. Biochem. Soc. Trans. 48, 2089–2100. 10.1042/BST20200241.

39. Hirota, T., Gerlich, D., Koch, B., Ellenberg, J., and Peters, J.-M. (2004). Distinct functions of condensin I and II in mitotic chromosome assembly. J. Cell Sci. 117, 6435–6445. 10.1242/jcs.01604.

40. Samejima, K., Booth, D.G., Ogawa, H., Paulson, J.R., Xie, L., Watson, C.A., Platani, M., Kanemaki, M.T., and Earnshaw, W.C. (2018). Functional analysis after rapid degradation of condensins and 3D-EM reveals chromatin volume is uncoupled from chromosome architecture in mitosis. J. Cell Sci. 131. 10.1242/jcs.210187.

41. George, C.M., Bozler, J., Nguyen, H.Q., and Bosco, G. (2014). Condensins are Required for Maintenance of Nuclear Architecture. Cells 3, 865–882. 10.3390/cells3030865.

42. Zhang, T., Paulson, J.R., Bakhrebah, M., Kim, J.H., Nowell, C., Kalitsis, P., and Hudson, D.F. (2016). Condensin I and II behaviour in interphase nuclei and cells undergoing premature chromosome condensation. Chromosome Res. 24, 243–269. 10.1007/s10577-016-9519-7.

43. Smits, V.A.J., and Gillespie, D.A. (2015). DNA damage control: regulation and functions of checkpoint kinase 1. FEBS J. 282, 3681–3692. 10.1111/febs.13387.

44. Toledo, L., Neelsen, K.J., and Lukas, J. (2017). Replication Catastrophe: When a Checkpoint Fails because of Exhaustion. Mol. Cell 66, 735–749. 10.1016/j.molcel.2017.05.001.

45. Hendriks, I.A., Lyon, D., Su, D., Skotte, N.H., Daniel, J.A., Jensen, L.J., and Nielsen, M.L. (2018). Site-specific characterization of endogenous SUMOylation across species and organs. Nat. Commun. 9, 2456. 10.1038/s41467-018-04957-4.

46. Cox, J., and Mann, M. (2008). MaxQuant enables high peptide identification rates, individualized p.p.b.-range mass accuracies and proteome-wide protein quantification. Nat. Biotechnol. 26, 1367–1372. 10.1038/nbt.1511.

47. Cox, J., Neuhauser, N., Michalski, A., Scheltema, R.A., Olsen, J.V., and Mann, M. (2011). Andromeda: a peptide search engine integrated into the MaxQuant environment. J. Proteome Res. 10, 1794–1805. 10.1021/pr101065j.

48. Cox, J., Hein, M.Y., Luber, C.A., Paron, I., Nagaraj, N., and Mann, M. (2014). Accurate proteome-wide label-free quantification by delayed normalization and maximal peptide ratio extraction, termed MaxLFQ. Mol. Cell. Proteomics 13, 2513–2526. 10.1074/mcp.M113.031591.

49. Tyanova, S., Temu, T., Sinitcyn, P., Carlson, A., Hein, M.Y., Geiger, T., Mann, M., and Cox, J. (2016). The Perseus computational platform for comprehensive analysis of (prote) omics data. Nat. Methods 13, 731–740.

50. Johnson, W.E., Li, C., and Rabinovic, A. (2007). Adjusting batch effects in microarray expression data using empirical Bayes methods. Biostatistics 8, 118–127. 10.1093/biostatistics/kxj037.

51. Perez-Riverol, Y., Bai, J., Bandla, C., García-Seisdedos, D., Hewapathirana, S., Kamatchinathan, S., Kundu, D.J., Prakash, A., Frericks-Zipper, A., Eisenacher, M., et al. (2022). The PRIDE database resources in 2022: a hub for mass spectrometry-based proteomics evidences. Nucleic Acids Res. 50, D543–D552. 10.1093/nar/gkab1038.

52. Ran, F.A., Hsu, P.D., Wright, J., Agarwala, V., Scott, D.A., and Zhang, F. (2013). Genome engineering using the CRISPR-Cas9 system. Nat. Protoc. 8, 2281–2308. 10.1038/nprot.2013.143.

53. Nielsen, C.F., Zhang, T., Barisic, M., Kalitsis, P., and Hudson, D.F. (2020). Topoisomerase IIα is essential for maintenance of mitotic chromosome structure. Proc. Natl. Acad. Sci. U. S. A. 117, 12131–12142. 10.1073/pnas.2001760117.

54. Meijering, A.E.C., Sarlós, K., Nielsen, C.F., Witt, H., Harju, J., Kerklingh, E., Haasnoot, G.H., Bizard, A.H., Heller, I., Broedersz, C.P., et al. (2022). Nonlinear mechanics of human mitotic chromosomes. Nature 605, 545–550. 10.1038/s41586-022-04666-5.

